# Carbon metabolism shapes FtsZ levels and cell division in a cyanobacterium

**DOI:** 10.1101/2025.05.26.656067

**Authors:** Wen-Shuo Ran, Xiaoli Zeng, Cheng-Cai Zhang

## Abstract

Cyanobacteria, as photoautotrophs, play key ecological roles and are widely used in synthetic biology research. While metabolism profoundly influences cellular processes like cell division, the regulatory mechanisms linking these pathways remain poorly understood in cyanobacteria. Here, we engineered the model cyanobacterium *Anabaena* sp. PCC 7120 by introducing an extra carboxylation module into the TCA pathway, perturbing this central metabolic pathway. This modification alters the division protein FtsZ levels, impairing cell division under varying light conditions. We found that 2-oxoglutarate, which is known as a metabolic signal, regulates *ftsZ* expression via the transcription factor NtcA. Furthermore, metabolic status modulates cell division in an NtcA-dependent manner, revealing a direct integration of metabolic control and cell division. Our findings uncover a coordination mechanism between metabolism and cell division in cyanobacteria, providing mechanistic insights for synthetic biology engineering and the understanding of metabolic regulation.

## Introduction

How different cell activities are orderly and effectively coordinated is a central question in biology. For a bacterial cell facing constantly changing environments, coordinating various metabolic and physiological processes is essential for survival and adaptation. Each bacterial species can maintain its distinct cell size and morphology by integrating the external environmental input and internal metabolic state ^1–5^. By integrating informational cues, bacterial cell division and the other cell cycle events, such as cell growth, DNA replication, segregation, etc., are thus coordinated ^6^. However, because of the complexity of such regulation, the mechanisms required are mostly poorly understood.

Bacterial cell division is a highly controlled, dynamic process involving numerous proteins. The prokaryotic tubulin homolog FtsZ is the central component for cell division in most bacteria, as well as plastids in algae and plants, and in some mitochondria ^6,7^. FtsZ can assemble into a ring-like structure called the Z-ring at the division site (mid-cell). The Z-ring then recruits the downstream proteins to form a multiprotein complex termed the divisome that drives cell constriction ^6–8^. Thus, the amount and the localization of the FtsZ protein in the cell must be tightly regulated. Changes in *ftsZ* expression level or FtsZ localization can strongly impact cytokinesis, which in turn can alter cell morphology, cell size, or even affect cell viability ^9–11^. Several factors regulating the activity or polymerization state of FtsZ have been reported ^12^; in contrast, much less is known about the mechanism that controls FtsZ levels in response to internal or external cues. Multiple promoters are found at the *ftsZ* genes in various bacteria ^13^, but few transcriptional regulators have been identified, and the underlying mechanism of the transcription regulation of *ftsZ* expression remains unclear. Recently, an increasing number of sRNAs, including cis– and trans-acting sRNAs, such as DinR, StfZ ^14^, DicF ^15^, and OxyS ^16^, have been found to regulate FtsZ levels post-transcriptionally. In addition, the amount of FtsZ levels was shown to be regulated by the ClpXP protease in different bacteria, such as *E. coli*^17^, *Bacillus subtilis* ^18^, *Caulobacter crescentus* ^19^, and *Synechocystis* sp. PCC6803 ^20^.

As one of the most fundamental biological processes, cell division is strongly linked to central carbon metabolism in different heterotrophic bacteria ^21,22^. In *E. coli* and *B. subtilis*, the accumulation of uridine diphosphate (UDP)-glucose stimulates the interaction between the glucosyltransferase OpgH/ UgtP and FtsZ, delaying the division machinery assembly, and hence, cell division ^23,24^. It was also reported that pyruvate mediates Z-ring localization and assembly in *B. subtilis* by regulating the localization of the pyruvate dehydrogenase ^25^. In *E. coli*, perturbations of acetyl-CoA metabolism impact cell size and division through changes in fatty acid synthesis. Moreover, other metabolic enzymes, such as glutamate dehydrogenase in *Caulobacter crescentus* and *Brucella abortus* ^26^, KidO (oxidoreductase) in *C. crescentus* ^27^, and *aceE*, *ackA*, *ccr*, *pta*, *tktA* in *E. coli* ^28^, have also been identified for their function in cell division. The absence of these metabolic enzymes causes cell division abnormalities to varying degrees, but their underlying mechanisms remain unknown.

Cyanobacteria are the only prokaryotes able to perform oxygenic photosynthesis, using light energy to convert water and CO_2_ into biomass ^29^. Its carbon metabolism is characterized by the Calvin-Benson cycle (carbon fixation) and photorespiration ^30^. Furthermore, cyanobacteria have an incomplete TCA cycle due to the lack of 2-oxoglutarate dehydrogenase (2-OGDH) which catalyzes succinyl-CoA production ^31^. Although some different bypass pathways were found to close the TCA cycle in different cyanobacteria species ^32,33^, biochemical analyses revealed that the contribution of these bypass pathways in terms of metabolic flux and metabolite pools to the TCA pathway is limited ^34^. A few studies suggest that the inorganic carbon and nitrogen regime ^3^ or nutrient metabolism can affect cell division in cyanobacteria ^35^, but the underlying regulation mechanism is unknown.

Since the relationship between metabolism and cell division remains unknown in cyanobacteria, we sought to disturb the TCA metabolic pathway by engineering a strain harboring an additional carboxylation module, using the model cyanobacterium *Anabaena* sp. PCC 7120 (hereafter *Anabaena*). This module contains the crotonyl-CoA carboxylase/reductase (Ccr), the most efficient carboxylase known in nature, to redirect metabolic flux into the TCA pathway. *Anabaena* is a filamentous cyanobacterium capable of diazotrophic growth following deprivation of combined nitrogen. Nitrogen fixation occurs exclusively in heterocysts, cells differentiated from vegetative cells within 20-24 h after nitrogen starvation ^36^. The process of heterocyst development is triggered by the accumulation of 2-oxoglutarate (2-OG), a signal of carbon-nitrogen metabolic balance ^36,37^. 2-OG then binds to NtcA to initiate the process of heterocyst differentiation ^37–40^. Our data reported here found that the addition of the extra module of carboxylation in *Anabaena* affects the pool of the TCA metabolites, 2-OG, which controls FtsZ levels through NtcA. This study highlights a distinct coordination mechanism between carbon metabolism and cell division in cyanobacteria, offering critical insights into the metabolic engineering of these photoautotrophic organisms.

## Results

### Introduction of an additional carboxylation module into *Anabaena*

One way to investigate the relationship between carbon metabolism and cell division is to disturb metabolic pools and examine their consequence on cell division. To do so, we integrated an extra carboxylation module (named PCEM), containing the genes *<u>p</u>co* (propionyl-CoA oxidase, <u>P</u>co), *ccr* (crotonyl-CoA carboxylase, <u>C</u>cr), *epi* (emC/mmC epimerase, <u>E</u>pi), and *mcm* (methyl malonyl-CoA isomerase, <u>M</u>cm) into the *Anabaena* genome with the substitution of the *alr2634-all2640* gene cluster (encoding peptide synthetases and polyketide synthases) by the Cpf1-based gene editing technique ^41^. The four genes of the module were in one artificial operon under the control of a previously characterized CT promoter (P_CT_, inducible by Cu^2+^ and theophylline) (Figs 1A and 1B) ^5^. Pco is the acyl-CoA oxidase ACX4 from *Arabidopsis thaliana* with a T134L mutation and a strong activity for oxidizing propionyl-CoA into acrylyl-CoA ^42^. Ccr is from *Methylobacterium extorques* and can produce methyl malonyl-CoA via reductive carboxylation of acrylyl-CoA with high efficiency ^42^. Mcm and Epi are from *Rhodobacter sphaeroides*, which together catalyze the methylmalonyl-CoA to succinyl-CoA ^42^. The sequential reactions catalyzed by these enzymes result in succinyl-CoA formation from propionyl-CoA, fixing one extra carbon (CO_2_) at the expense of one NADPH. This CO_2_ fixation pathway provides a direct metabolic link to the incomplete TCA cycle in *Anabaena* (Fig. 1A and S1 Fig). The resulting cyanobacterial strain (gPCEM) was verified by PCR as shown in Fig. 1C. Further western blotting experiments showed that each of the 4 genes was expressed in the gPCEM strain in BG11 48 h after the addition of 1 mM theophylline and 0.3 µM Cu^2+^ (determined as the optimal inducer concentrations, which will be used in the following studies unless otherwise indicated) with continuous standard light (SL) illumination (Fig 1D). These results indicated that the PCEM module could be well expressed in *Anabaena*.

**Fig 1.**
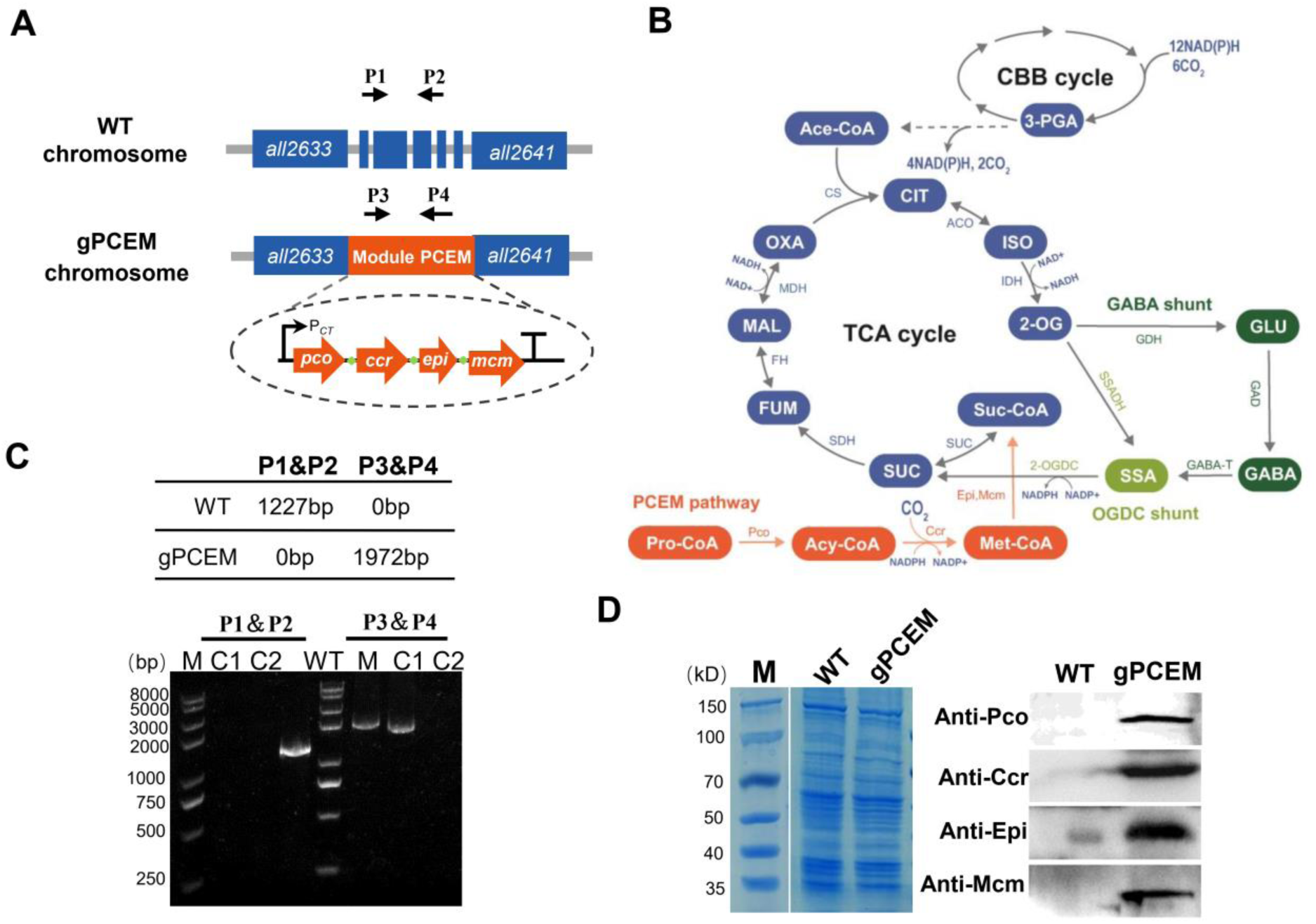
Engineering the PCEM carboxylation module in *Anabaena*. (A) Schematic presentation for the construction of the gPCEM strain. The promoter and terminator for the PCEM module are indicated by a polyline arrow and a “T” shape, respectively. Green dots indicate the ribosome binding sites of each gene. The position of the primers for genotype verification by PCR in (C) was shown as a short arrow. P1, P2, P3, and P4 are the short names for the primers Pall2637-F, Pall2639-R, PccrF-22, and PmcmR-22, respectively (Table S3). Genes of the PCEM module are colored in orange, and genes from the *Anabaena* genome are colored in blue. *pco*: the gene encoding the propionyl-CoA oxidase, *ccr*: the gene encoding the crotonyl-CoA carboxylase, *epi*: the gene encoding the emC/mmC epimerase, *mcm*: the gene encoding the methyl malonyl-CoA isomerase ^42^. (B) An overview of the PCEM module in metabolic pathways and its intersection with the TCA pathway. Cyanobacteria lack the 2-OGDH (2-oxoglutarate dehydrogenase) enzyme in the TCA pathway. The OGDC shunt that converts 2-OG to succinate by SSADH (succinic semialdehyde dehydrogenase) and 2-OGDC (2-oxoglutarate decarboxylase) is shown in light green, and the GAGB shunt that converts 2-OG to succinate via GLU and GABA is indicated in dark green. The PCEM module that converts propionyl-CoA to succinyl-CoA is shown in orange. 3-PGA: 3-Phosphoglyceric acid; Ace-CoA: acetyl CoA; CIT: citrate; ISO: isocitrate; 2-OG, 2-oxoglutarate; Suc-CoA: succinyl-CoA; SUC: succinate; FUM: fumarate; MAL: malate; OXA: oxaloacetate; ACO, aconitase; CS, citrate synthase; FH, fumarate hydratase; GABA, g-amino butyric acid; GAD, glutamate decarboxylase; GDH, glutamate dehydrogenase; GLS, glutaminase; ICL; isocitrate lyase; IDH, isocitrate dehydrogenase; 2-OGDH, 2-oxoglutarate dehydrogenase; LPS, lipopolysaccharide; MD, malate dehydrogenase; MS, malate synthase; OAA, oxaloacetate; PDH, pyruvate dehydrogenase; SUC, succinyl-CoA synthetase; MDH, malate dehydrogenase; SSA, succinic semialdehyde; SSADH, SSA dehydrogenase. CBB cycle: Calvin–Benson–Bassham cycle. TCA cycle, tricarboxylic acid cycle. (C) PCR verification of the gPCEM strain. The expected sizes of the PCR products for WT and gPCEM strains are shown in the upper part of the panel. C1: colony 1, C2: colony 2 (D) Western blotting analysis of the levels of the indicated proteins in the indicated strains. Similar amounts of total proteins extracted from different strains were loaded on the gel, stained with Coomassie Brilliant Blue (left, CBB), or probed with polyclonal antibodies against Pco, Ccr, Epi, or Mcm (right).

### Metabolic pool analysis of gPCEM strain under different light conditions

How does this extra carboxylation module affect the TCA metabolite pools in *Anabaena*? To answer this question, the products of the PCEM module, and the TCA pathway, including propionyl-CoA, pyruvate, succinate, fumarate, malate, citrate, 2-OG, and acetyl-CoA of the gPCEM strain at different light conditions (HL, high light; SL, standard light; LL, low light, see details in Materials and Methods) were analyzed by Liquid Chromatography-Mass Spectrometry (LC-MS) ^43^. As shown in Figure 2, propionyl-CoA, the first substrate of the PCEM module, exhibited a dramatic reduction in the gPCEM strain under different light conditions. These results indicated that the heterologous Pco has strong catalytic activity in *Anabaena* (Fig. 2). On the contrary, metabolites of the TCA pathway in the gPCEM strain varied significantly under different light conditions (Fig. 2). Under LL, the intracellular levels of fumarate, malate, and 2-OG in the gPCEM strain exhibited significant reduction, whereas the amounts of succinate and citrate remained relatively stable when compared to the WT. However, under SL and HL, the levels of most of the TCA cycle metabolites in the gPCEM strain increased significantly, as compared to the WT strain, such as succinate, fumarate, malate under SL, and fumarate, malate, citrate, 2-OG under HL. These results suggested that introducing the PCEM module into *Anabaena* could cause different disturbances to the metabolic pools of the TCA pathway under different light conditions. Notably, pyruvate, the end product of the glycolysis pathway, and acetyl-CoA, a central metabolic intermediate, remained in the gPCEM strain similar to those in the WT strain across all light conditions except for pyruvate which showed a reduction in the gPCEM strain under low light (LL).

**Fig 2.**
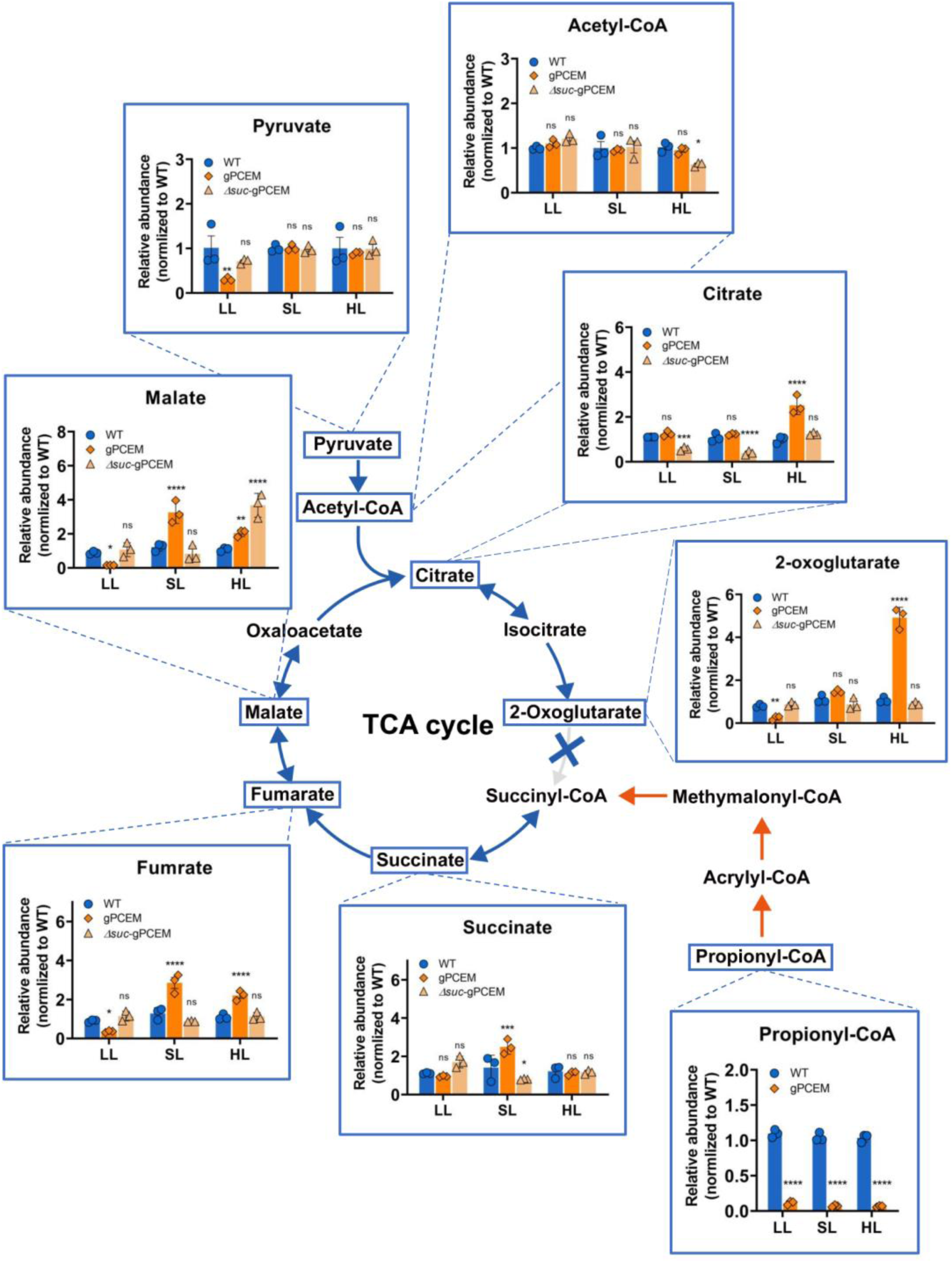
Changes of the selected metabolite pools in the gPCEM under different light conditions. Graphical representation of relative changes of the metabolites between the WT, the gPCEM, and the Δ*suc*-gPCEM at different light conditions, respectively. *Δsuc-gPCEM* strain: the *sucC* and *sucD* genes were deleted together (Δ*suc*) in the gPCEM strain background. SucC and SucD are succinyl-CoA synthetases converting succinyl-CoA to succinate. The data presented here indicate results from three biological replicates. The student’s t-test examined the differences between the WT and the other strains. Results were considered significant at * p < 0.1, ** p < 0.01, *** p < 0.001, **** p < 0.0001, ns: no significance. LL: low light; SL: standard light; HL: high light.

### The presence of the PCEM module affects cell differentiation in *Anabaena*

One intermediate from the TCA pathway, 2-OG, acts as a nitrogen-carbon balancing signal in cyanobacteria ^36,37^. In *Anabaena*, the accumulation of 2-OG signals nitrogen starvation and triggers heterocyst differentiation ^36–39^. Since 2-OG levels in the gPCEM strain under different light conditions were changed significantly, we expected that heterocyst development would be impacted. Therefore, we examined their effect on heterocyst development in BG11_0_, a culture medium free of combined nitrogen. The WT strain was analyzed in parallel as a control. When observed under a microscope, the gPCEM strain under different light conditions was affected in heterocyst frequency (S2A Fig). After 48 h of growth in BG11_0_ medium, the gPCEM strain could form mature heterocysts under all three different light conditions; however, the percentage of heterocysts showed a decrease under LL (6% ± 0.31%), an increase under SL (11.7% ± 0.62 %) and under HL (15.6% ± 0.55%), as compared to the WT at each of the corresponding conditions (7.2% ± 0.32%; 9.4% ± 0.32%, 9.4% ± 0.56%) (S2A Fig). Moreover, a Mch (multiple-contiguous-heterocysts) phenotype was occasionally observed under SL and HL (S2A Fig). To confirm this phenotype, we further quantified the pattern of heterocysts along the filaments. The heterocyst intervals of the WT strain showed regularity under different light conditions, with 18 to 20 vegetative cells, 12 to 13 vegetative cells, and 9 to 11 vegetative cells as the most representative intervals under LL, SL, and HL, respectively. In the gPCEM strain, heterocyst distributions kept a regularity under LL, but the intervals increased to 24 to 25 vegetative cells, consistent with a lower heterocyst frequency. Under SL and HL, heterocyst patterns of the gPCEM strain were changed, since a high frequency of Mch was observed and the intervals were shortened to 9 to 11 vegetative cells and 5 to 7 vegetative cells as the most representative intervals under SL and HL, respectively (S2B Fig). These results are consistent with the changes in the 2-OG levels observed in the gPCEM strain under different light conditions, further confirming the metabolite disturbances caused by the PCEM module as measured by the LC-MS/MS.

### The presence of the PCEM module affects the cell morphology of *Anabaena*

To evaluate the effects following the metabolic disturbances of the TCA pathway caused by the PCEM module in *Anabaena*, we first tested cell growth under different light conditions in BG11 medium in the presence of inducers as described above. As shown in Figure 3A, the growth rates of the gPCEM strain were slightly slower than that of the wild type under different light conditions, and the difference increased with increasing light intensities (Fig. 3A). When observed under a microscope, the gPCEM strain exhibited significant cell morphology defects under LL and HL, and the phenotype became more pronounced with the extension of culture time, while under SL conditions, its cell morphology was comparable to that of the WT (Figs 3B and 3C). Under LL, over time, the average cell length of the gPCEM strain gradually increased while the cell width remained constant, suggesting a delay in cell division (Fig. 3C). On the 6th day, the average cell length of the gPCEM strain was 9.3 µm as compared to 3.6 µm found for cells of the WT strain (Fig. 3C). Under HL, the cell morphology of the gPCEM strain becomes more and more irregular over time, as both cell length and width gradually increase with a large variation (Figs 3B and 3C). As a control, we confirmed that the markless deletion of the *all2634-all2640* gene cluster did not cause any cell morphology defects in *Anabaena* under similar conditions (S3A Fig), consistent with previous reports ^41^. These results all suggested that the expression of the extra carboxylation module may affect cell growth and cell division in *Anabaena*.

**Fig 3.**
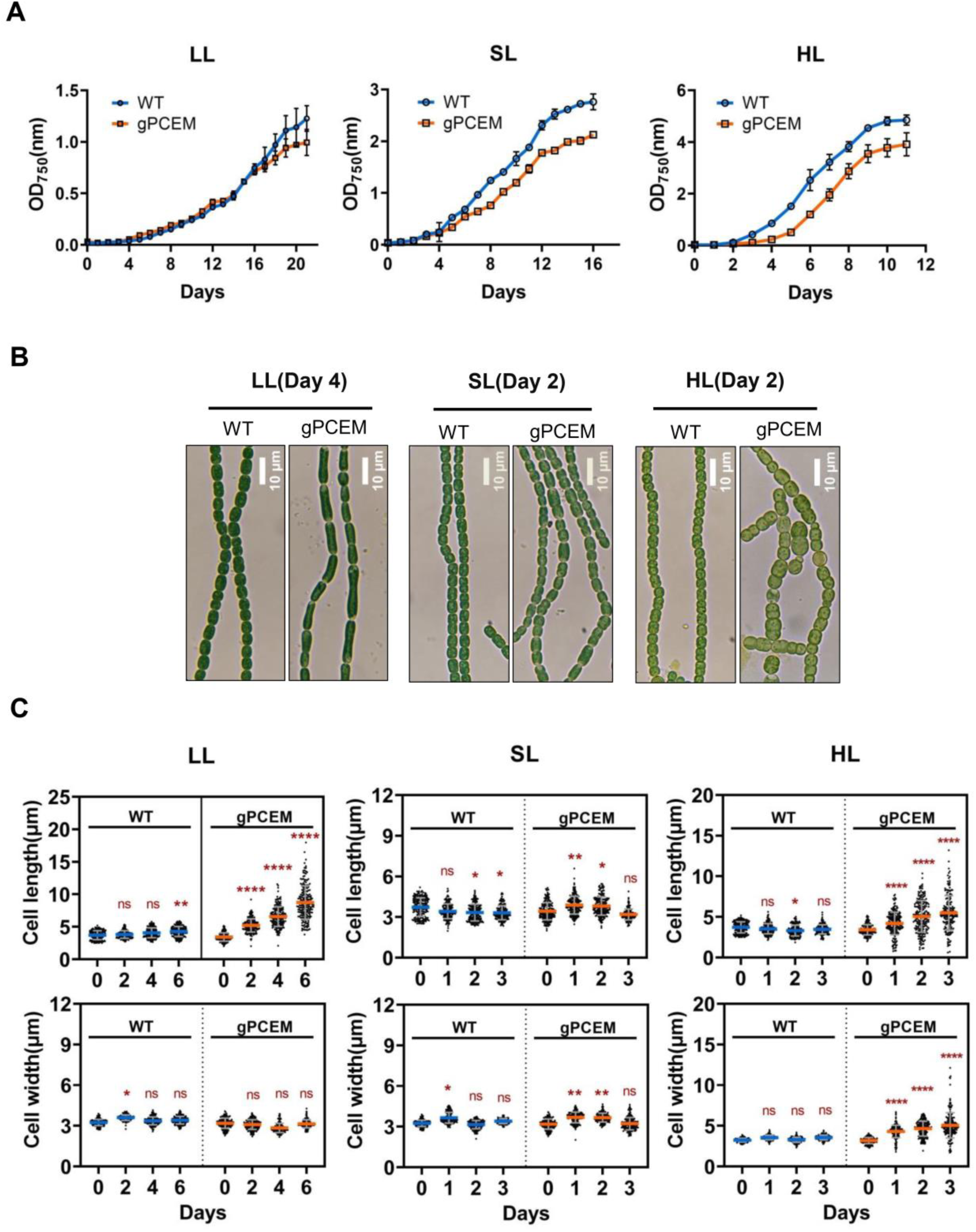
Phenotype of the gPCEM strain. (A) The growth curves of the WT and gPCEM strain in BG11 medium in the presence of 1 mM of theophylline (TP) and 0.3 μM Cu^2+^ under indicated light conditions. LL: low light; SL: standard light; HL: high light. The initial concentration measured at OD_750_ for each strain was 0.05. All values are shown as mean ± standard deviation, calculated from triplicate data. (B) Micrographs of *Anabaena* filaments of the WT (wild type) and the gPCEM strain in the indicated days and light conditions. Scale bars: 10 µm. (C) Statistical analysis of cell length and width of the WT (wild type) and the gPCEM strain at the indicated days after induction under light conditions. 300 cells of each strain from three independent experiments were measured. The student’s t-test examined the differences between the 0 days and the other days. Results were considered significant at * p < 0.1, ** p < 0.01, *** p < 0.001, **** p < 0.0001, ns: no significance.

To rule out potential side effects from the introduction of multiple genes as the cause of the observed cell division and morphological changes, we constructed a series of strains related to the PCEM module in the WT background, including four strains expressing each of the four single genes (strain WT::OE_CT_-*pco*, WT::OE_CT_-*ccr*, WT::OE_CT_-*epi,* WT::OE_CT_-*mcm*), and three strains expressing two, three, or all fours genes of the whole module (strain WT::OE_CT_-*PC* with the *pco-ccr* genes, strain WT::OE_CT_-*PCE* with the *pco-ccr-epi* triple genes, and strain WT::OE_CT_-*PCEM* with all four genes of the PCEM module). All these constructs were carried on a replicative plasmid, and gene expression was under the control of the CT promoter, inducible by copper and theophylline. After conjugation, we successfully obtained colonies for all strains, except WT::OE_CT_-*PCEM*. Finally, a few colonies of WT::OE_CT_-*PCEM*, validated by PCR, could be obtained only when a trace amount of copper was removed from glassware with acid treatment. These colonies grew poorly on plates without inducers. However, when inoculated in liquid BG11 medium, the strain WT::OE_CT_-*PCEM* was unable to grow, in contrast to other strains and the WT. We further observed the filament of the strain WT::OE_CT_-*PCEM* from the colonies under a microscope and found that its cells displayed significant cell morphology defects, much stronger than those of the gPCEM strain under HL (S3B Fig). These observations suggest that even without inducers, basal-level expression of the whole module derived from a high-copy plasmid leads to cell growth defects. The other strains were further inoculated in BG11 medium with 1 mM theophylline and 0.3 µM Cu^2+^ under SL. Western blotting results confirmed that Pco, Ccr, Mcm, and Epi proteins were present in the strains containing the corresponding genes (S3C Fig). When observed under a microscope, the cell morphology of all these strains was comparable to that of the WT (S3D Fig). Similar results were obtained when these strains were grown under LL. Together, these results indicate that the metabolic disturbances and phenotypic changes in the gPCEM strain arise from the integrated activity of the entire PCEM module, rather than from the individual expression or function of its constituent enzymes.

### *ΔsucCD* suppresses cell morphology defects in the gPCEM strain

The final product of the carboxylation PCEM module is presumed to be succinyl-CoA (Fig. 1B), which could potentially enter and disrupt the TCA pathway (Fig. 1B). While the 2-OGDH (2-oxoglutarate dehydrogenase) is absent in cyanobacteria ^31^, previous studies in *Synechocystis* sp. PCC 6803 ^32^ and *Synechococcus* sp. PCC 7002 ^34^ has shown that succinyl-CoA is still converted to succinate via succinyl-CoA synthetase (encoded by *sucC* and *sucD*). Our sequence alignment analysis confirmed the presence of SucC and SucD homologs in *Anabaena* (S4 Fig), supporting the possibility that PCEM-generated succinyl-CoA may enter the TCA pathway via succinate, potentially causing metabolite disturbances and, hence, cell morphology defects in the gPCEM strain. To test this hypothesis, we deleted the *sucC* and *sucD* genes together (*Δsuc*) in the gPCEM strain backgrounds, leading to strain *Δsuc*-gPCEM. We first checked the cell morphology of this strain under different light conditions (Fig. 4). While the gPCEM strain exhibited significant cell morphology changes under LL and HL, these phenotypes were largely suppressed in the *Δsuc*-gPCEM strain (Fig. 4A). The average cell length and width of the *Δsuc*-gPCEM strain are similar to those of the WT, independent of light conditions (Fig. 4B). Furthermore, heterocysts in *Δsuc*-gPCEM also exhibit similar frequencies and distribution patterns as the WT strain (S2 Fig). LC-MS metabolite profiling showed that while the propionyl-CoA was reduced similarly to the gPCEM strain, most of the TCA pathway metabolites in the Δ*suc*-gPCEM strain showed similar variation patterns as the WT strain under comparable light conditions (Fig.2). We did observe changes in the levels of a few metabolites under certain conditions, including acetyl-CoA (slight reduction under HL), citrate (a decrease under LL and SL), succinate (a slight reduction under SL), and malate (a significant increase under SL) (Fig 2). Importantly, these changes did not cause a particular phenotype in cell division and morphology in this strain. These observations suggest that SucC/D-mediated metabolism plays a central role in the morphological abnormalities of the gPCEM strain.

**Fig 4.**
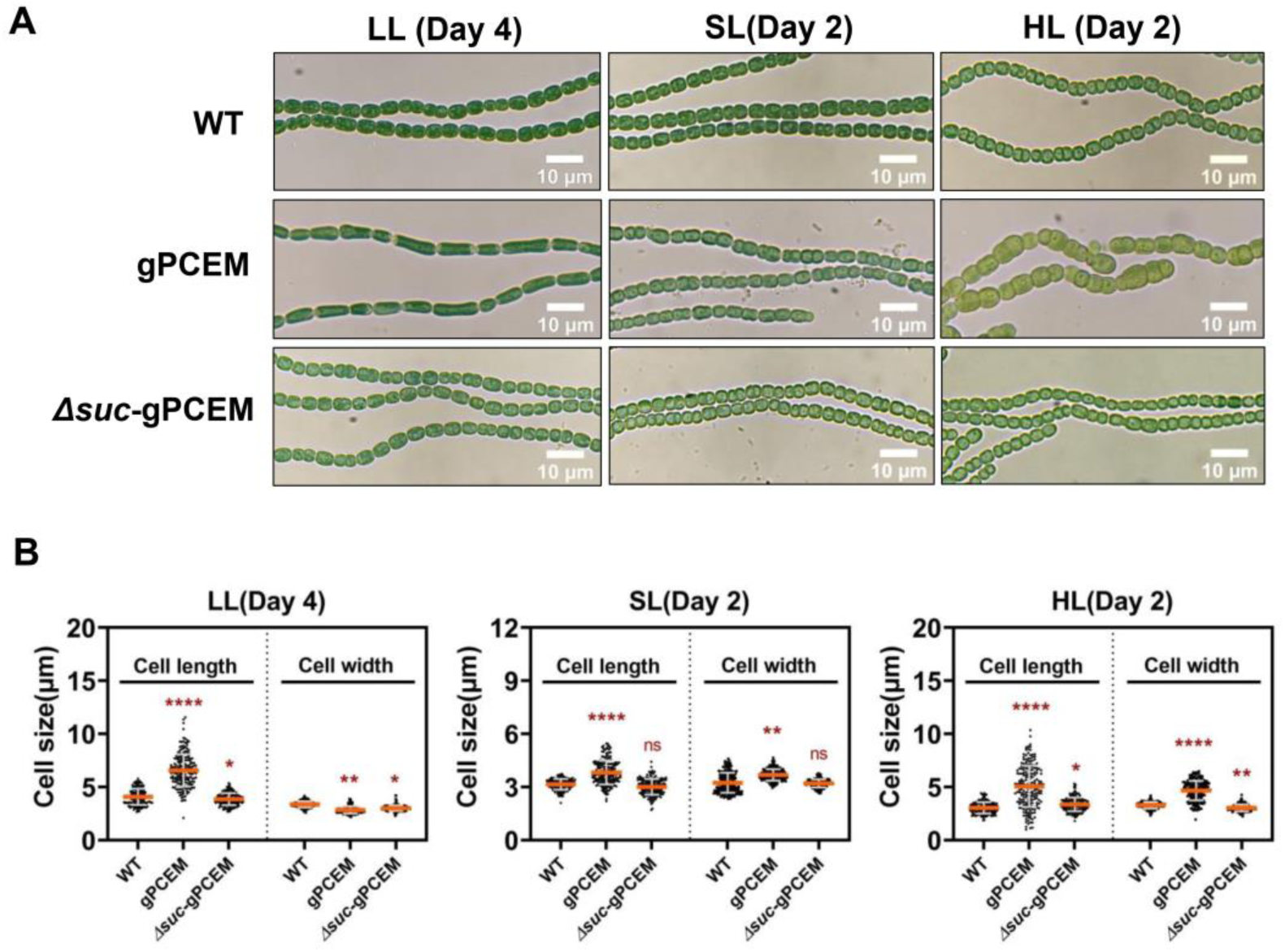
***ΔsucCD* suppresses the cell morphology defects of the gPCEM strain.** (A) Micrographs of *Anabaena* filaments of the WT, the gPCEM strain, and the Δ*suc*-gPCEM strain cultured in BG11 medium with 1 mM of theophylline (TP) and 0.3 μM Cu^2+^ at the indicated days and light conditions. LL: low light; SL: standard light; HL: high light. Scale bars: 10 µm. (B) Statistical analysis of cell length and cell width of the indicated strains at the indicated days after induction under different light conditions. 300 cells of each strain from three independent experiments were measured. The student’s t-test examined the differences between the WT and the other strains. Results were considered significant at * p < 0.1, ** p < 0.01, *** p < 0.001, **** p < 0.0001, ns: no significance.

### The gPCEM strain has different FtsZ levels in vivo under different light conditions

Given that in bacteria, FtsZ dosage critically regulates cell division, with reduced levels causing filamentation and excess levels triggering minicell formation ^9,11^, we hypothesized therefore the expression of the PCEM module under different light conditions may affect the FtsZ levels in the *gPCEM* strain, and cause the corresponding cell morphology defects. To validate this hypothesis, the intracellular FtsZ levels of the gPCEM strain under different light conditions were tested in comparison to the WT under similar conditions. As shown in Fig. 5A, under SL, the FtsZ levels of the gPCEM and the WT strains remained the same, independent of the incubation time (Fig. 5A). However, the FtsZ levels of the gPCEM strain decreased significantly under LL over time but increased significantly under HL over time (Fig. 5A). These results indicated that the cell morphology defects of the gPCEM strain under LL and HL are correlated with the changes in the FtsZ levels. We further analyzed the FtsZ levels of the *Δsuc*-gPCEM strain at different light conditions (Fig. 5B). As expected, they all showed FtsZ levels similar to those of the WT strain, independent of the light conditions. These results suggested that the changes in FtsZ levels in the gPCEM strain under LL and HL require the connection between the PCEM module and the TCA pathway.

**Fig 5.**
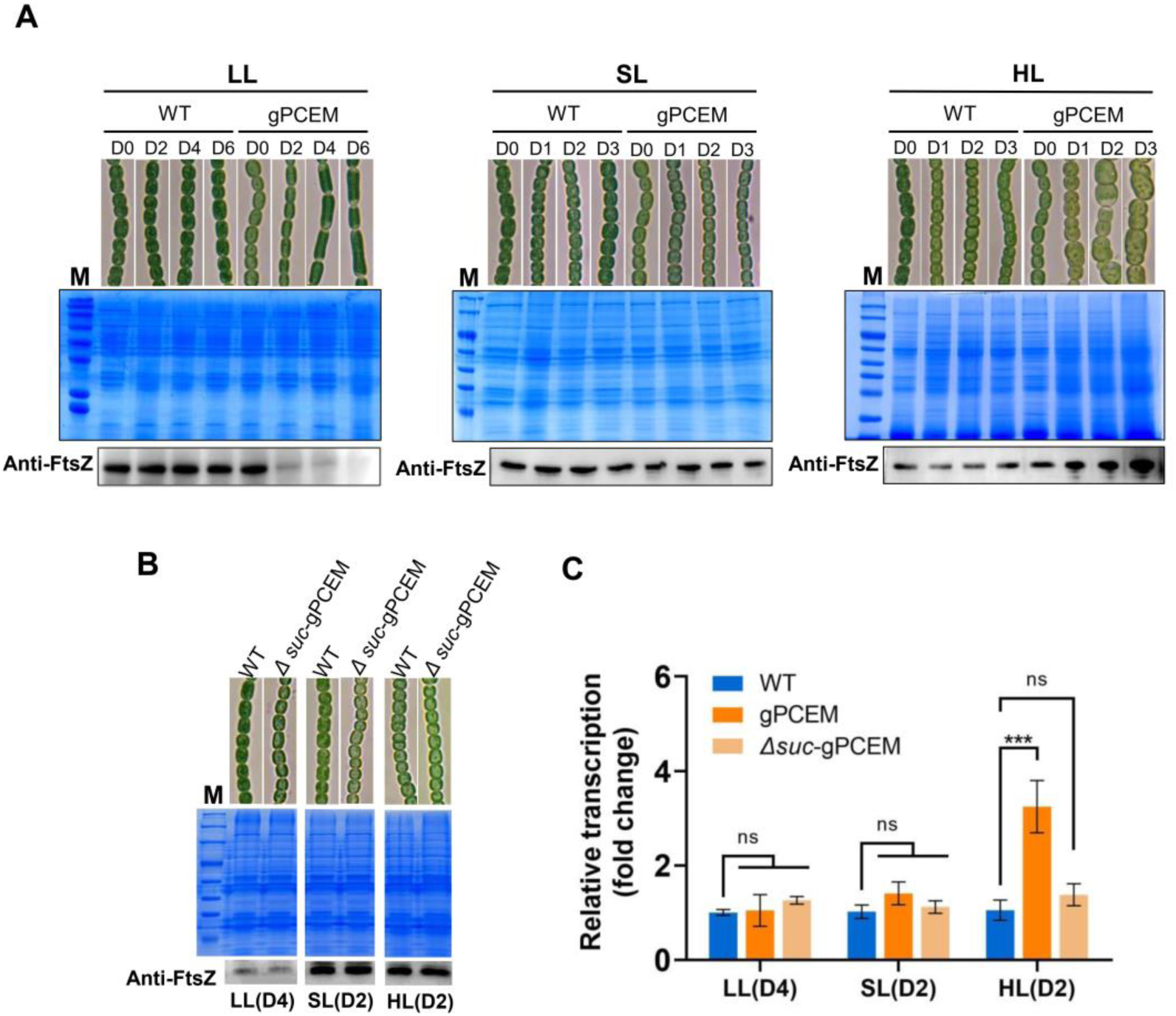
The FtsZ levels in the gPCEM strain vary under different light conditions. (A) Western blotting of FtsZ levels in the WT and the gPCEM strain cultured in BG11 medium with 1 mM of theophylline (TP) and 0.3 μM Cu^2+^ at different days and light conditions. Similar amounts of total proteins extracted from different strains were loaded on the gel, stained with coomassie brilliant blue (middle, CBB), or probed with polyclonal antibody against FtsZ (Bottom). Representative microscopic images of the indicated strains at different conditions are presented. LL: low light; SL: standard light; HL: high light. (B) FtsZ levels of the Δ*suc*-gPCEM strain in comparison with those of the WT and the gPCEM strain at the indicated days and light conditions as in (A). D0-D6: day 0 to day 6. (C) The transcription levels of the *ftsZ* gene in the WT, the gPCEM strain, and the Δ*suc*-gPCEM strain at the indicated time points and light conditions. The experiment was done with three technical replicates in three biological replicates. The student’s t-test examined the differences between the WT and the other strains under the same conditions. Results were considered significant at ***p < 0.001, *p < 0.05. ns: no significance.

To evaluate whether the transcriptional levels of *ftsZ* also change in the gPCEM strain under LL and HL, quantitative Real-Time PCR (qRT-PCR) was performed using the gPCEM, the *Δsuc*-gPCEM, and the WT strains. Under different tested conditions, the transcript levels of *ftsZ* remained relatively stable in all strains, except for the gPCEM strain cultured under HL, which showed a significant increase, consistent with the increase in FtsZ protein levels under similar conditions (Fig. 5C). These results indicate that transcriptional regulation of *ftsZ* expression contributes to the increase of FtsZ protein levels in the presence of the PCEM module under HL. However, under LL, the decrease in the FtsZ levels observed in the gPCEM strain was likely caused by posttranscriptional regulations that remain to be determined.

### NtcA regulates the transcription of the *ftsZ* gene in *Anabaena*

How do the TCA metabolite pools affect the transcriptional level of *ftsZ*? Under HL conditions, the gPCEM strain exhibits elevated levels of fumarate, malate, citrate, and 2-oxoglutarate (2-OG) (Fig.2). Since fumarate and malate levels are also significantly elevated in the gPCEM strain under SL conditions (Fig. 2), while FtsZ levels remain unchanged (Fig.5), the elevated levels of 2-OG and citrate are likely the main factors for the enhanced transcriptional activity of the *ftsZ* gene under HL conditions. Previous studies indicated that the 2-OG signal in *Anabaena* could be perceived by the global transcription factor NtcA, which regulates the transcription of genes involved in different processes ^36–40^, and could regulate the expression of the *mre* genes (encoding MreB, MreC, and MreD, cell elongation) related to cell division ^3^. The promoters of the NtcA regulon usually contain the consensus palindromic sequence GTAN8TAC at the upstream of the transcription start site (TSS) ^40^. We wondered whether the *ftsZ* gene could also be regulated by NtcA in *Anabaena*. We first checked the promoter region of *ftsZ* for the presence of potential NtcA binding motifs by FIMO (https://meme-suite.org/meme/tools/fimo). Interestingly, two putative NtcA-binding sites (GTAN8TAT and GTN10AC, –142bp and –470bp to the start codon) at the *ftsZ* promoter region could be identified (*p*-value < 0.001, Fig 6A). According to the published transcriptomic data ^44^, six TSS sites could be found at the *ftsZ* promoter region, and the two putative NtcA-binding sites were located in front of these two TSSs (Fig 6A). To confirm that NtcA could recognize the two putative NtcA-binding sites, electrophoretic mobility shift assays (EMSA) were performed. Two PCR-amplified DNA fragments of 200 bp from the *ftsZ* promoter region, each containing one of the two putative binding sites, respectively, were obtained. As shown in Figures 6B and 6C, NtcA alone could not bind to the two tested DNA fragments, even with increasing amounts of NtcA (Figs 6B and 6C). However, with the presence of 2-OG, both DNA fragments formed a DNA-protein complex with NtcA even at a low protein concentration (Figs 6B and 6C). The addition of increasing concentrations of 2-OG significantly promoted the formation of the NtcA-DNA complex (Figs 6B and 6C). These results indicated that NtcA could bind to the promoter region of *ftsZ* in response to the 2-OG signal in vitro.

**Fig 6.**
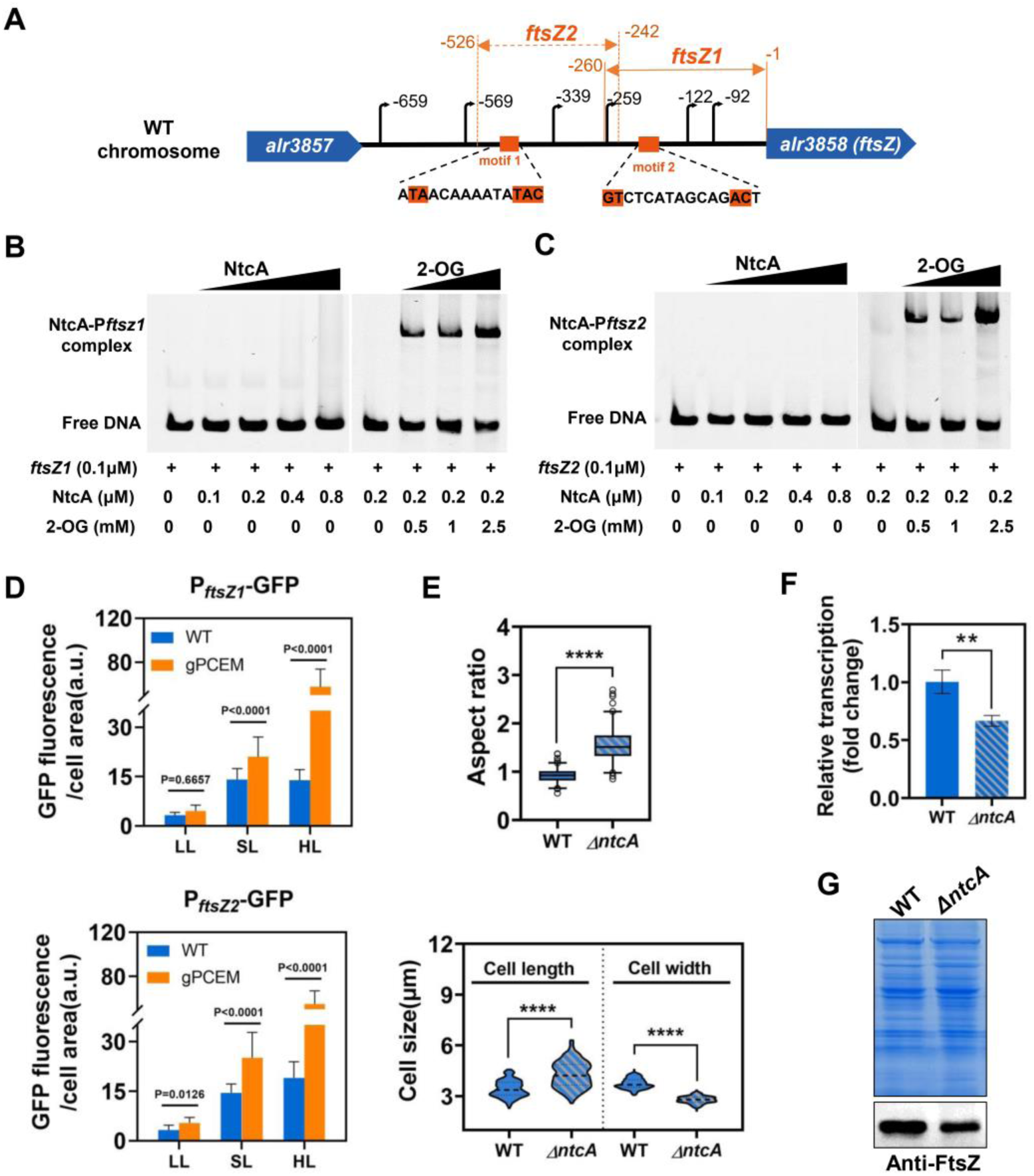
NtcA binds to the promoter region of the *ftsZ* gene in the presence of 2-OG (2-oxoglutarate). (A) Diagram of the positions of the TSS sites of *ftsZ* (indicated by polyline arrow), the putative NtcA binding motifs (in orange) at the promoter region of *ftsZ*, and the DNA fragments (indicated as *ftsZ1* and *ftsZ2*) used for the assays. (B) EMSA assays demonstrate the binding of NtcA to the two putative motifs, *ftsZ1* (B) or *ftsZ2* (C) in the presence of 2-OG. In both Figure B and C, the Left panel, two FAM-labeled DNA, *ftsZ1* or *ftsZ2* (2 pmol), were first incubated with increasing concentrations of the NtcA protein. Right panel, 2 pmol DNA fragments were incubated with 4 pmol NtcA in the absence or the presence of increasing concentrations of 2-OG. (D) Statistical analysis of GFP fluorescence in the WT and the gPCEM strains that had a *gfp* gene under the control of the *ftsZ1* (–260bp to –1bp) or the *ftsZ2* (–526bp to –242bp) promoter at the indicated light conditions, based on images as shown in Figure S5 and S6. 100 cells of each strain from three independent experiments were measured. (E)Statistical analysis of aspect ratio, cell length, and cell width of cells of WT *Anabaena* and the *ntcA* mutant strains grown in BG11_0_ with 5 mM NH4^+^ under standard light conditions for 24 hours. The aspect ratio was calculated by dividing the length of the axis parallel to the filament by the length of the axis perpendicular to the filament. 200 cells of each strain from three independent experiments were measured. The student’s t-test examined the differences between the WT and the *ntcA* mutant. Results were considered significant at * p < 0.1, ** p < 0.01, *** p < 0.001, **** p < 0.0001, ns: no significance. (F) The transcription levels of the *ftsZ* gene in the WT *Anabaena* and the *ntcA* mutant grown in BG11_0_ with 5 mM NH^4+^ under standard light conditions at 24 hours. ** p < 0.01.(G) Western blotting of FtsZ in the WT *Anabaena* and the *ntcA* mutant grown in BG11_0_ with 5 mM NH4^+^ under SL conditions at 24 hours.

To investigate whether the two newly identified NtcA motifs are responsible for the transcriptional regulation of *ftsZ* in vivo, the promoter regions, each containing one NtcA motif, were fused to *gfp*. The fusions carried on a replicative plasmid were introduced into the WT and the gPCEM strains, respectively. The GFP fluorescence from these fusions in different strains under different light conditions was determined. Under LL, no significant GFP fluorescence could be detected for all tested strains (Figs 6D and S5, and S6 Figs). Under SL, the GFP fluorescence from both fusions in the gPCEM strain was a little bit brighter than that of the WT strains, suggesting a transcriptional activation of *ftsZ* under such conditions in the WT is similar to the gPCEM strain (Fig. 6D and S5 and S6 Figs). Under HL, the GFP fluorescence from both fusions in the gPCEM strain was much brighter than that of the WT strain (Fig. 6D and S5 and S6 Figs). These results are consistent with the changes in the transcriptional levels of *ftsZ* analyzed by qRT-PCR above, and the increase of the 2-OG levels under similar conditions in gPCEM (Fig. 5C). They also indicated that the two promoter regions were involved in the *ftsZ* transcriptional regulation in vivo.

### NtcA mutant displays morphological defects

Following the finding that elevated 2-OG levels in gPCEM affect cell morphology through NtcA, we wondered whether *ntcA* controls *ftsZ* expression under physiological conditions in the wild type. In our analysis of cell length, cell width, FtsZ levels, and 2-OG levels in the gPCEM strain under varying light conditions, we used the WT as a control for a similar analysis. We observed that the average cell length of WT *Anabaena* gradually decreased as light intensity increased, while cell width remained largely unchanged (S7A Fig). This phenotype correlated with elevated levels of 2-OG and increased expression of FtsZ (S7B-D Figs). Although the 2-OG levels in WT *Anabaena* showed no significant change from SL to HL, we noted that the increased NtcA protein levels under HL led to further enhancement of FtsZ expression (S7D Fig). These findings provide additional evidence for NtcA-mediated FtsZ regulation in *Anabaena*.

If NtcA regulates *ftsZ* expression, we expect that a *ntcA* mutant would display a cell-division-related phenotype. A previous study already reported that the *ntcA* mutant (CSE2) in ammonium-supplemented BG11_0_ medium exhibits a higher cell aspect ratio than the wild-type strain ^3^. NtcA regulates the expression of the *mre* operon, and the Mre proteins control peptidoglycan synthesis along the bacterial lateral cell wall, hence cell elongation; therefore, the phenotype of aspect ratio was proposed to be caused by elevated levels of the Mre proteins in the *ntcA* mutant ^45^. Our findings corroborate this observation, as we also detected a higher cell aspect ratio in the *ntcA* mutant under the same conditions (Fig. 6E), with a significant increase in cell length and a reduction in cell width (Fig. 6E). Further analysis revealed that both the level of *ftsZ* transcripts and that of the FtsZ protein were significantly elevated in the *ntcA* mutant (Figs 6F-G). These results indicate that NtcA regulates the expression of FtsZ in *Anabaena*. Taking all results together, we conclude that NtcA regulates the transcription of the *ftsZ* gene in *Anabaena* in a 2-OG-dependent manner. The observed morphological abnormalities in the *ntcA* mutant can be attributed to a synthetic effect caused by decreased FtsZ levels and increased Mre levels, which together account for changes in the aspect ratio.

## Discussion

In heterotrophic bacteria, central carbon metabolism—including the glycolytic pathway, the pentose phosphate pathway, and the TCA cycle—is closely linked to cell division ^22,24,25,27,28^. The molecular mechanisms by which different carbon metabolites coordinate the regulation of cell division vary significantly, highlighting the complexity of the network between carbon metabolism and cell division. Despite the significant differences in metabolic modes as compared to heterotrophic bacteria, cyanobacteria also modulate cell size and morphology according to nutrient regimes (such as carbon supply) and light intensity ^2,3,45^; however, the mechanism remains largely unexplored. In this study, we disturbed the functioning of the TCA pathway in the model cyanobacterium *Anabaena* by introducing a highly efficient, artificial CO_2_ fixation module (PCEM module) (Figs 1, 2, and 4). We observed changes in the levels of FtsZ (Fig. 5) as well as cell morphology (Figs 3 and 4). Combining genetics and biochemistry techniques, we confirmed that the TCA cycle intermediate 2-oxoglutarate (2-OG) and the transcription factor NtcA play a crucial role in coordinating cell division with growth and carbon metabolism in Cyanobacteria (Fig. 6), which provides a distinct mechanism in contrast to model heterotrophic bacteria. The PCEM module could convert propionyl-CoA to succinyl-CoA, fixing one extra carbon (CO_2_) and consuming one additional NADPH in the process (Fig. 1) ^42^. Since the succinyl-CoA can integrate into the TCA pathway via succinyl-CoA synthetase (SucC and SucD) in cyanobacteria (Figs 2 and 4) ^34^, the gPCEM strain may redirect propionyl-CoA flux via the engineered pathway. While the observed cell division defects in the gPCEM strain could theoretically result from depletion of propionyl-CoA precursors-which remain uncharacterized in cyanobacteria – three lines of experimental evidence argue against this possibility: Firstly, control strains expressing individual PCEM components (the WT::OE_CT_-*pco* strain, the WT::OE_CT_-*PC* strain, as well as the WT::OE_CT_-*PCE* strain), all exhibited normal cell division (S3 Fig), excluding nonspecific effects of protein expression or potential propionyl-CoA consumption. Secondly, genetic disruption of the metabolic steps connecting the PCEM-TCA cycle in the *Δsuc-gPCEM* completely suppressed the cell division phenotype (Figs 4 and 5), demonstrating the involvement of a functional metabolic coupling in the alteration of cell division. Finally, we measured the acetyl-CoA levels under different conditions in the gPCEM strain and found no evidence of significant alteration of this metabolite (Fig. 2). The only condition where a drop in acetyl-CoA levels was observed was under high light (HL) in the *Δsuc*-gPCEM strain, where decreased acetyl-CoA showed no phenotypic consequence.

The PCEM module displays varying efficiency in carbon source input in *Anabaena* under various light conditions, affecting the TCA cycle differently (Fig. 2). This indicates that the PCEM-TCA coupled metabolic pathway in the gPCEM strain is light-dependent. This effect may be explained by the effect of light intensity on the production of NADPH in Cyanobacteria ^46,47^, which is a reducing power required for CO_2_ fixation by PCEM. High light intensity may enhance the production of NADPH through photosynthesis in *Anabaena*, conditions in favor of the PCEM-TCA coupled metabolism. In contrast, low light intensity limits the energy supply and substrates available for the PCEM-TCA coupled pathway, ultimately affecting its overall activity. In addition, the TCA pathway may be directly or indirectly regulated by light so that carbon metabolism operates in response to photosynthetic activities. On the other hand, the artificial and synthetic PCEM pathway itself is unlikely to be regulated by light, consistent with the observation that consumption of propinyl-CoA (Fig. 2), the first substrate of this pathway, is similar under different light conditions tested.

Based on the impact of the PCEM module on the TCA pathway (Fig. 2), we found that FtsZ levels are controlled through distinct mechanisms under different light conditions in *Anabaena* (Figs. 5, 6, and 7). Under LL conditions, the gPCEM strain exhibits low levels of several metabolites and the FtsZ protein compared to the WT (Fig. 2). However, the transcriptional levels of the *ftsZ* gene remain comparable to those in the WT (Fig. 5C). This observation suggests that these TCA metabolites may directly or indirectly influence the FtsZ amount at posttranscriptional levels and Z-ring assembly through an unknown mechanism. Consistent with these results, posttranscriptional control of *ftsZ* expression^14–16^, or FtsZ degradation ^17–19^, has been suggested previously, although the underlying mechanism remains unknown.

Our studies also confirm a mechanism that accounts for the relationship between the TCA metabolite 2-OG levels and the transcriptional control of *ftsZ* (Fig. 7C). Combining the results of the EMSA assay, the P_ftsz_-*gfp* reporter expression analyses, and the *ntcA* mutant analyses, we proposed that the global transcription factor NtcA regulates the transcription of the *ftsZ* gene in a 2-OG-dependent manner in *Anabaena* (Figs. 6 and 7C). The accumulation of 2-OG and the NtcA protein level in the gPCEM strain under highlight led to an increased transcript level of *ftsZ,* resulting in irregular cell division (Figs 2 and 7C, S7 Fig). High levels of FtsZ will initiate FtsZ ring formation in an uncontrolled manner, causing ectopic cell division sites, and thus the formation of cells with varying sizes and shapes (Fig. 3). Such a phenotype is similar to that observed in *E. coli* when FtsZ levels increase with mini cell formation ^9^. In the WT *Anabaena*, we observed a significant reduction in cell length (indicating higher cell division rates) with increasing light intensity. This correlated with a rise in the intracellular 2-OG levels from low light (LL) to standard light (SL) or high light (HL), as well as a consistent increase in NtcA protein levels in cells shifted from LL to HL. Based on these findings, we propose that varying light intensities lead to differential intracellular 2-OG levels, which in turn modulate NtcA-dependent cell division (Figs 7A and 7B). However, the physiological advantages of this light-dependent morphological regulation remain unclear and warrant further investigation. Furthermore, the presence of multiple TSS, including two NtcA-recognized promoters (Fig. 6), suggests that the expression of *ftsZ* is under complex control with the possibility of integrating various environmental inputs. Notably, citrate-a key metabolite in the TCA pathway that accumulates alongside 2-OG in the gPCEM strain under HL-may also contribute to the regulation of *ftsZ* expression, though this possibility requires further validation.

**Fig 7.**
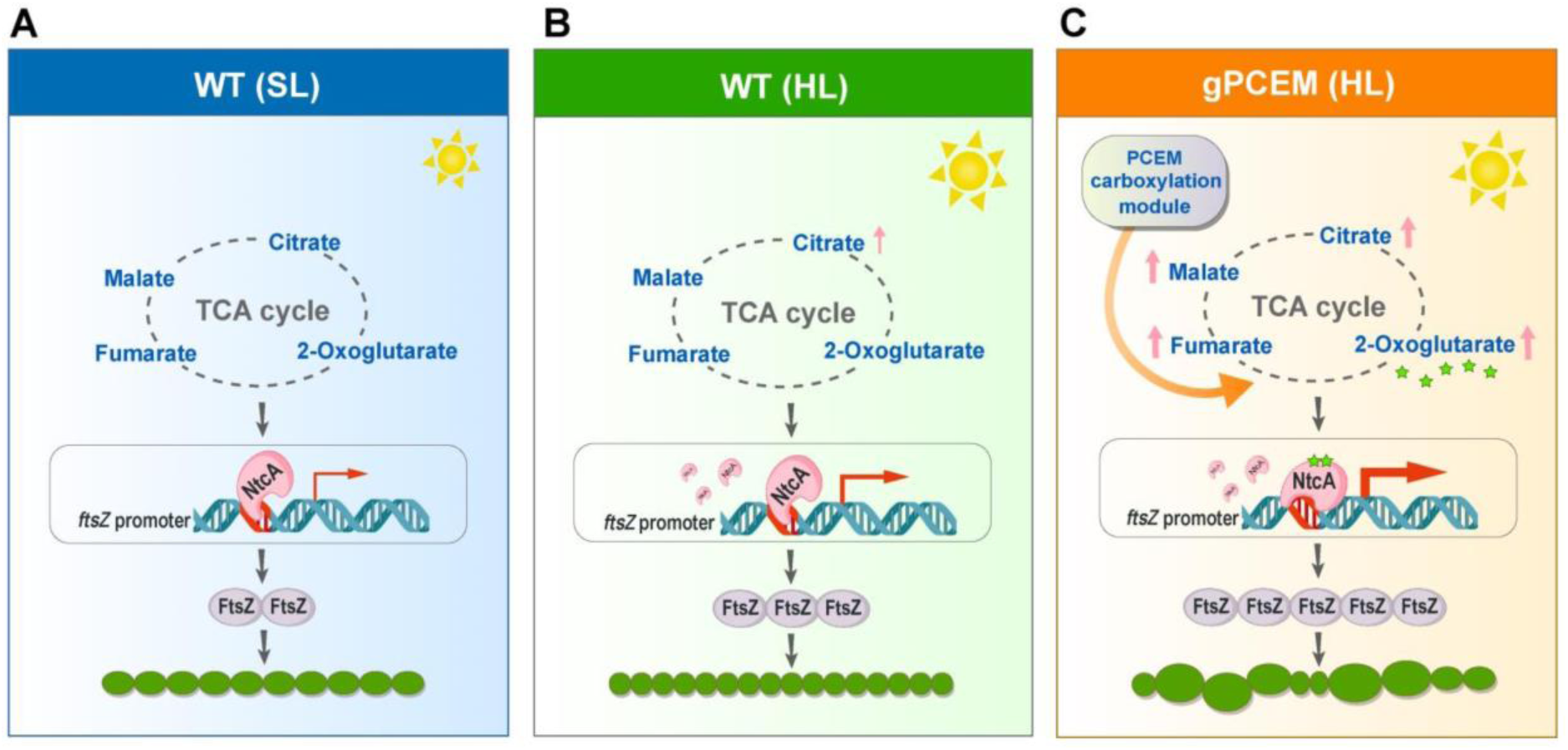
**A model for NtcA-regulated FtsZ levels and cell division in *Anabaena.*** A: Under SL (standard light) conditions, the TCA pathway in the WT operates at normal efficiency, leading to no significant changes in carbon metabolites. NtcA regulates FtsZ expression at a basal level, and cell division occurs normally without noticeable changes in cell morphology. B: Under HL (high light) conditions, the efficiency of the TCA pathway slightly increased, resulting in elevated levels of succinate and citrate in the WT *Anabaena* strain compared to SL conditions. This metabolic shift enhances the NtcA protein level, which in turn increases FtsZ expression. The elevated FtsZ levels promote faster cell division, resulting in reduced cell length. C: Under HL conditions, the TCA pathway in the gPCEM strain exhibits high efficiency, causing significantly elevated levels of fumarate, malate, citrate, and 2-OG (2-oxoglutarate, indicated by a green star). The accumulation of NtcA and 2-OG plays a major role in increasing FtsZ levels by enhancing the transcription of the *ftsZ* gene through the transcription factor NtcA. Consequently, the elevated FtsZ level promotes asymmetric cell division, leading to irregular cell morphology.

Previous studies have shown that NtcA negatively regulates the expression of the *mre* genes (encoding MreB, MreC, and MreD) ^3^. FtsZ controls cell width by regulating peptidoglycan synthesis at the division site, whereas MreB controls peptidoglycan synthesis at the lateral cell wall ^8,10^. Since both *mre* and *ftsZ* are under the regulation of NtcA, the observed cell morphological defects in the gPCEM strain could be the consequence of the misregulation of both systems. These findings suggest that NtcA, as a global transcription factor, coordinates cell division and elongation in response to C-to-N balance in cyanobacteria. 2-OG accumulates transiently and serves as a signal at the early phase of heterocyst differentiation. Whether 2-OG/NtcA coordinately regulates cell division during the developmental process remains an open question.

## Methods

### Strains and Cultural Conditions

The strains used in this study are listed in Table S1. *Anabaena* WT and its derivatives were cultivated at 30°C in BG11 medium with continuous illumination (standard light, 30 μmol/m^2^ s). When needed, 1 mM of theophylline (TP) and 0.3 μM Cu^2+^ were added to induce gene expression under indicated light conditions (LL, low light, 7 μmol/m^2^ s; SL, standard light, 30 μmol/m^2^ s; HL, high light, 70 μmol/m^2^ s). For heterocyst induction, cells grown to the logarithmic phase in BG11 were transferred to BG11_0_ (BG11 without combined nitrogen) medium, and heterocysts were observed under a light microscope. To make the data comparable, TP and Cu^2+^ were added to both the WT and the gPCEM strains before and during heterocyst induction. To compare the growth of the WT and the gPCEM strains, the absorbance at 750 nm was measured at the indicated time points after inoculation in BG11 liquid medium with TP and Cu^2+^ under indicated light conditions. The *ntcA* mutant, as described in ^41^, was first cultured at 30 °C in BG11 medium with 5 mM NH4^+^ under SL to log phase and then transferred to BG11_0_ medium with 5 mM NH4^+^ under SL for further analysis.

### Construction of Plasmids and Cyanobacterial Recombinant Strain

All plasmids and primers used in this study are listed in Tables S2 and S3, respectively. All *Anabaena* variants, including the Δ*alr2634-all2640* strain, the gPCEM stain, and the *Δsuc*-gPCEM strain were generated by the genome editing technique based on CRISPR-Cpf1, and the plasmids were constructed as previously described ^41^. The plasmids for gene overexpression in *Anabaena* were constructed based on the PCT vector, allowing control of gene expression by the artificial CT promoter ^4,5^. The plasmids for P_ftsZ1/2_-*gfp* transcription fusion in *Anabaena* WT or in the gPCEM stain were constructed with a modified PCT vector in which the CT promoter elements were deleted so that the fusion is only under the control of the native *ftsZ* promoter ^4,5^. To generate recombinant strains or mutants, the corresponding plasmid was transferred into *Anabaena* by conjugation as described ^4^.

### Microscopy

Cells were collected by filtration and then broken with FastPrep-24 (6.0 m/s, 60 s) in a protein-loading buffer as described ^5^. Cell extracts were heated at 100°C for 5-10 min, followed by centrifugation at 12000 rpm for 5 min. The supernatant was collected and separated on 10% sodium dodecyl sulfate-polyacrylamide gel electrophoresis and then transferred to the nitrocellulose membranes. Blots were probed with the corresponding protein rabbit antiserum (1:1000) and HRP-conjugated secondary antibody (1:5000, goat anti-rabbit).

### Quantitative Real-Time PCR

The WT, gPCEM, and *Δsuc*-gPCEM strains were cultivated in BG11 with 1 mM of theophylline (TP) and 0.3μM Cu^2+^ under the indicated light conditions. Cultures after 4 days under LL, and 2 days under SL and HL were harvested by rapid filtration. The total RNA was extracted using a Plant Total RNA Isolation Kit (#RE-05011, Foregene, China), and the genomic DNA was removed by using the RNase-free DNase I (Promega, USA). The obtained total RNA was reverse transcribed using the HiScript®II Q RT SuperMix (#R223-0, Vazyme, China). Quantitative Real-Time PCR was performed by using ChamQ SYBR qPCR Master Mix (Vazyme). The housekeeping gene *rnpB* was used as an internal control. The data were analyzed by ABI 7500 SDS software. The transcriptional levels of the *ftsZ* gene were quantified with the comparative CT method (2-ΔΔCT method). Primers used for qPCR are listed in Table S3.

### Electrophoretic Mobility Shift Assays (EMSA)

The His-tagged NtcA protein (NtcA-His6) for EMSA was prepared as previously described ^39^. The DNA fragments were amplified from the genome of *Anabaena* using the primers Palr3858F260m / Palr3858R1 m and Palr3858F526m / Palr3858R242m, respectively (Table S3). The two DNA fragments were further labeled with fluorescent 6-carboxyfluorescein (FAM) tag at 5’ end. 2 pmol of the labeled probes were incubated with varying amounts of NtcA-His6 and 2-OG in a binding buffer (50 mM Tris-HCl pH7.5, 60 mM KCl, 5 mM MgCl_2_, 0.25 M NaCl, 0.5 mM EDTA, 5 mM Beta-mercaptoethanol, 12% glycerol,40 μg/mL BSA) for 30 min at 25°C. The samples were subjected to a 5% polyacrylamide gel electrophoresis and run in 1X PAGE buffer (50 mM Tris-HCl pH 8.0, 2 mM EDTA, 380 mM glycine) at constant 20 mA for 3 hours in an ice bath. Imaging and data analysis were performed using Amersham Typhon.

### Extraction and quantification of cellular Metabolites by LC-MS/MS

The WT *Anabaena*, the gPCEM stain, and the *Δsuc*-gPCEM strain, cultured as indicated above, were harvested by rapid filtration and resuspended in 500 μL of ice-cold extraction buffer (80% methanol). After incubating at –20 °C for 20 min, the cells were broken by ultrasonication at 90W amplitude for 1 min. The samples were further incubated at –20 °C for 15 min and then centrifuged at 13,000 rpm for 10 min at 4 °C. The supernatant was transferred to a new 1.5 mL tube, evaporated, and dried by freeze-vacuum drying (Eppendorf Concentrator Plus).

To quantify the organic acids in the samples, the pellets were resuspended in 200 μL of the initial mobile phase (1:9 =solvent A: B). All samples were filtered with a 0.22 µm Ultra free-MC membrane before being subjected to the column. 1 μL of each sample was analyzed by LC-MS/MS on an Agilent 6550 iFunnel Q-TOF system, equipped with an Atlantis Premier BEH Z-HILIC column ^43^. The samples were separated at 40°C with a flow rate of 0.3 mL/min. The chromatography process was used as the following: gradient of solvent A (20 mM ammonium formate pH = 9.0) to solvent B (methanol): t = 1.0 min, A-10%: B-90%; t = 5.0 min, A-40%: B-60%; t = 6.0 min, A-70%: B-30%, t = 8.0 min, A-70%: B-30%; t = 8.5 min, A-10%: B-90%; t = 13 min (end of gradient). MS data were scanned over a range of 50-500 m/z at a scan rate of 1 spectra/sec. The parameters for mass spectrum were as follows: capillary voltage, 3.5 kV; nitrogen atomization (35 psig), drying (8 L/min, 200°C), and sheath gas (12 L/min, 400°C).

To quantify the CoA esters in the samples, the pellets were resuspended in 200 μL of the initial mobile phase (9.5:0.5 = solvent A: B). All samples were filtered with a 0.22 µm Ultra free-MC membrane before being subjected to the column. 20 μL of each sample was analyzed by LC-MS/MS on an Agilent 6550 iFunnel Q-TOF system, equipped with a 50 x 2.1 mm C18 column (Kinetex 1.7µm EVO C18 100 Å). The samples were separated at 25°C with a flow rate of 0.2 mL/min. The chromatography process was used as follows: gradient of solvent A (50 mM ammonium formate pH = 8.0) to solvent B (methanol): t = 2.0 min, A-95%: B-5%; t = 10.0 min, A-80%: B-20%; t = 12.0 min, A-80%: B-20%; t = 13.0 min, A-20%: B-80%; t = 15.0 min, A-20%: B-80%; t =15.1min, A-95%: B-5%; t = 19.0 min (end of gradient). MS data were scanned over a range of 80-1000 m/z at a scan rate of 1 spectra/sec. The parameters for mass spectrum were as follows: capillary voltage, 3.5 kV; nitrogen atomization (35 psig), drying (8 L/min, 320°C), and sheath gas (12 L/min, 400°C).

The corresponding standard solutions prepared at 50 μg/mL concentration in the initial mobile phase were analyzed in parallel. All standard stock solutions were prepared daily just before running samples in the LC-MS/MS system.

All LC/MS/MS peak data were collected on an Agilent MassHunter Data Acquisition software, and analyzed using Qualitative Analysis Navigator B.08.00 and Quantitative Analysis B.09.00. The signal intensity of each metabolite was integrated by the corresponding peak area and normalized to the WT strain under the same conditions. The differences (P < 0.05, indicating statistical significance) between the WT and the other strains were evaluated using the Student’s t-test.

## Acknowledgments

This research was supported by the CAS Project for Young Scientists in Basic Research (Grant No: #YSBR-015), the Strategic Priority Research Program of the Chinese Academy of Sciences (Grant No. XDB0480000), the Key Program of the National Natural Science Foundation of China (Grant No. 32330003) and the Youth Innovation Promotion Association CAS (Grant No. 2022342). We thank Amel Latifi (Aix-Marseille University) for providing the *ntcA* mutant strain.

## Author contributions

Conceptualization, CZ.Z., and X.Z.; Methodology, WS.R.; Investigation, WS.R.; Formal Analysis, WS.R., and X.Z.; Visualization, WS.R. and X.Z.; Writing – Original Draft: X.Z., Writing – Review & Editing: X.Z. and CC.Z.; Funding Acquisition: CC.Z. and X.Z.

